# Localization of TRP channels in healthy oral mucosa from human donors

**DOI:** 10.1101/2021.06.02.446798

**Authors:** Yalda Moayedi, Stephanie Michlig, Mark Park, Alia Koch, Ellen A Lumpkin

## Abstract

The oral cavity is exposed to a remarkable range of noxious and innocuous conditions, including temperature fluctuations, mechanical forces, inflammation and environmental and endogenous chemicals. How such changes in the oral environment are sensed by oral cells and tissues is not completely understood. Transient receptor potential (TRP) ion channels are a diverse family of molecular receptors that are activated by chemicals, temperature changes, and tissue damage. In non-neuronal cells, TRP channels play roles in inflammation, as well as tissue development and maintenance. In somatosensory neurons, TRP channels mediate nociception, thermosensation and chemosensation. To assess whether TRP channels might be involved in environmental sensing in the human oral cavity, we investigated the distribution of TRP channels in human tongue and hard palate. Oral biopsies were collected from volunteers and underwent fluorescent immunohistochemistry followed by confocal imaging. We analyzed immunoreactivity of TRP channels in human oral epithelia including TRPV3, TRPV4, TRPV1, TRPM8, and TRPA1. TRPV3 and TRPV4 were expressed in epithelial cells with inverse expression patterns where they are likely to contribute to epithelial development and integrity. TRPA1 immunoreactivity was found in fibroblasts, subsets immune cells, and neurons, consistent with known roles of TRPA1 in sensory transduction, as well as in response to damage and inflammation. TRPM8 immunoreactivity was found in lamina propria cells and some neuronal subpopulations including some neurons within the end bulbs of Krause, consistent with a role in thermal sensation. TRPV1 immunoreactivity was identified in intraepithelial nerve fibers, in some end bulbs of Krause, and in epithelial cells, consistent with roles in nociception and thermosensation. Immunoreactivity of TRPM8 and TRPV1 in end bulbs of Krause suggest that these structures contain a variety of neuronal afferents, including those that mediate nociception, thermosensation and mechanotransduction. Collectively, these studies support the role of TRP channels in oral environmental surveillance and response.

## Introduction

Oral mucosa are poised to transduce chemosensory and somatosensory stimuli during feeding, speech, and protection against biological and chemical agents. Sensory transduction occurs through the activation of receptor molecules that detect mechanical, chemical, or thermal stimulation of tissues. The Transient Receptor Potential (TRP) family of cation channels include molecular receptors that encode chemical, thermal, and mechanical aspects of environmental stimuli, and are important transducers of chemesthetic stimuli. TRPV1 is a heat-activated receptor that is also the molecular target of capsaicin, the pungent component of spicy chilies (Caterina et al. 1997). TRPV3 and TRPV4 are activated by warm temperatures, chemical stimuli, and osmotic swelling (Chung et al. 2004a; Peier et al. 2002b; Vogt-Eisele et al. 2007; Vriens et al. 2004). TRPM8 is activated by cooling, menthol, and other chemicals that produce a cooling sensation (McKemy et al. 2002; Peier et al. 2002a). TRPA1, the ‘wasabi receptor’, is a promiscuous damage sensory that is activated by noxious pungent compounds in radishes, mustard, and garlic, as well as reactive oxygen species produced during tissue stress (Bautista et al. 2005; Takahashi et al. 2008). In rodents, these TRP channels are expressed in somatosensory neurons; however, several are also reported to be expressed in epithelial cells, including the oral epithelium (Vandewauw et al. 2013; Wang et al. 2011).

With regard to oral functions, several TRP channels are important in flavor perception and pathophysiology. TRPM5 and TRPM4 are expressed in rodent and human type II taste cells and are essential components of the signaling pathways downstream of sweet, bitter, and umami stimuli (Dutta Banik et al. 2018; Liu and Liman 2003; Perez et al. 2002; Prawitt et al. 2003; Zhang et al. 2003). Similarly, in rodents TRP channels PKD1L3 and PKD2L1 are expressed in type III taste cells, although their contribution to taste-cell physiology is still debated (Horio et al. 2011; Ishimaru et al. 2006; Ye et al. 2016). TRPV4 has been found to regulate type III taste-cell differentiation in mice, loss of which results in reduced sensitivity to sour compounds (Matsumoto et al. 2019). In addition to roles in gustation, TRP channels are essential contributors to oral temperature transduction, chemesthesis, and response to injury. For example, TRPM8, TRPA1 and TRPV1 mediate sensory transduction of pungent chemicals in numerous flavor-enhancing spices, including mint, radishes, chiles, black pepper and cinnamon (Roper 2014). These TRP channels are also important for thermal transduction in the oral cavity (Lemon 2021). Furthermore, TRP channel expression and activation has been linked to oral cancer cell proliferation and pain in rodents and human cancers (Fujii et al. 2020; Okamoto et al. 2012; Ruparel et al. 2015).

Despite the importance of TRP channels for oral functions, the localization of these channels in healthy tissues from human oral cavity is not clear. In this study, we present an immunohistochemical analysis of TRP channels in human hard palates and tongue biopsies from healthy tissues.

## Methods

### Study enrollment criteria

Human studies were approved by the Institutional Review Board of Columbia University. Enrollment was performed as described in (Moayedi et al. 2021). Oral biopsies were collected from adult volunteers (27-45 years old n=12, **Supplementary Table 1**). Exclusion criteria: infection, pain, oral injury or cutaneous abnormality that could interfere with safety or data interpretation; anticoagulants (e.g. aspirin, coumadin, NSAIDs), bleeding disorder, keloidal or hypertrophic scarring history, oral cancer, neurological diseases, epithelial innervation abnormalities in biopsy site, known or suspected medical or psychological conditions that may affect ability to consent or to follow instructions for wound care, active medical conditions that may affect risk of infection or healing after biopsy.

### Informed consent

Written informed consent was obtained by study personnel prior to any protocol-specific procedures. The study was conducted in accordance with the Food and Drug Administration (FDA) approved revision of the Declaration of Helsinki, current FDA regulations, and International Conference on Harmonization guidelines.

### Tissue collection

Oral tissue collection was performed as described (Moayedi et al. 2021). Biopsies of either front of tongue or palate rugae were collected from each participant (**Supplementary Figure 1**). Biopsy site was anesthetized (2% lidocaine with epinephrine (1:100,000)). A 4-mm punch biopsy oriented perpendicular to the specimen and punch was taken down to the submucosal layer. College pliers were used to remove the core and reveal the submucosal layer and scissors used to free the biopsy if needed. The specimen was removed and placed in phosphate buffered saline (PBS) and pressure applied to the biopsy site. The biopsy was sutured closed if necessary and additional gauze applied. Compensation was given after biopsies were collected. Discarded human foreskin tissue was used to test antibody concentrations and for peptide blocking experiments.

### Tissue analysis

Tissues were embedded (TissueTech OCT), flash frozen, and 25 µm sections were made on gelatinized slides. Slides were incubated for 30 min at 37°C, fixed with 4% paraformaldehyde (0-15 min) followed by five washes in PBS. Slides were blocked in PBS with 0.1% triton-X 100 (PBST) and 5% normal goat serum. Sections were incubated overnight with primary antibodies (**Supplementary Table 2)** mixed in blocking buffer at 4°C. Slides were washed 3x in PBST and incubated with secondary antibody in blocking solution for 1–2 h, then then washed 5x in PBS and mounted in Fluoromount-G with DAPI. Specimens were imaged with a laser scanning confocal microscope equipped with 40X (NA 1.3) and 20X (NA 0.8) lenses. Antibody concentrations and staining parameters were optimized on foreskin tissue. In antigen blocking experiments, primary antibody cocktail was preincubated with 5x protein antigen concentration relative to antibody concentration 30 min at room temperature prior to incubation with sections. Results of antigen retrieval experiments are shown in **Supplementary Figure 2**. For each biomarker, 2–4 independent samples from front of tongue and hard palate rugae were tested.

## Results

To directly compare immunoreactivity of a panel of TRP channels in human oral cavity, biopsies of hard palate rugae or tongue papillae were collected from healthy adult volunteers (**Supplementary Table 1)**. Antibodies against TRP channel targets were first optimized on human foreskin tissue, and then tested on at least two independent samples of both tongue and hard palate rugae for each probe. When antigens were available, antigen blocking experiments were performed to test the specificity of the antibody to the antigen (**Supplementary figure 2**).

Expression of TRPV3 and TRPV4, two channels that show high expression in epithelial cells, was examined. Antigen blocking experiments showed specificity of TRPV3 antigen to this antibody (**Supplementary figure 2**). In oral tissue, TRPV3 immunoreactivity was found primarily in the basal epithelial layers of both hard palate and tongue mucosa (**Figure 1**). In the hard palate, TRPV3 localization in the basal epithelium overlapped with regions where Merkel cells are typically found (**Figure 1A)** (Moayedi et al. 2021). TRPV3 immunoreactivity was undetectable in oral neurons in this study.

**Figure 1.**
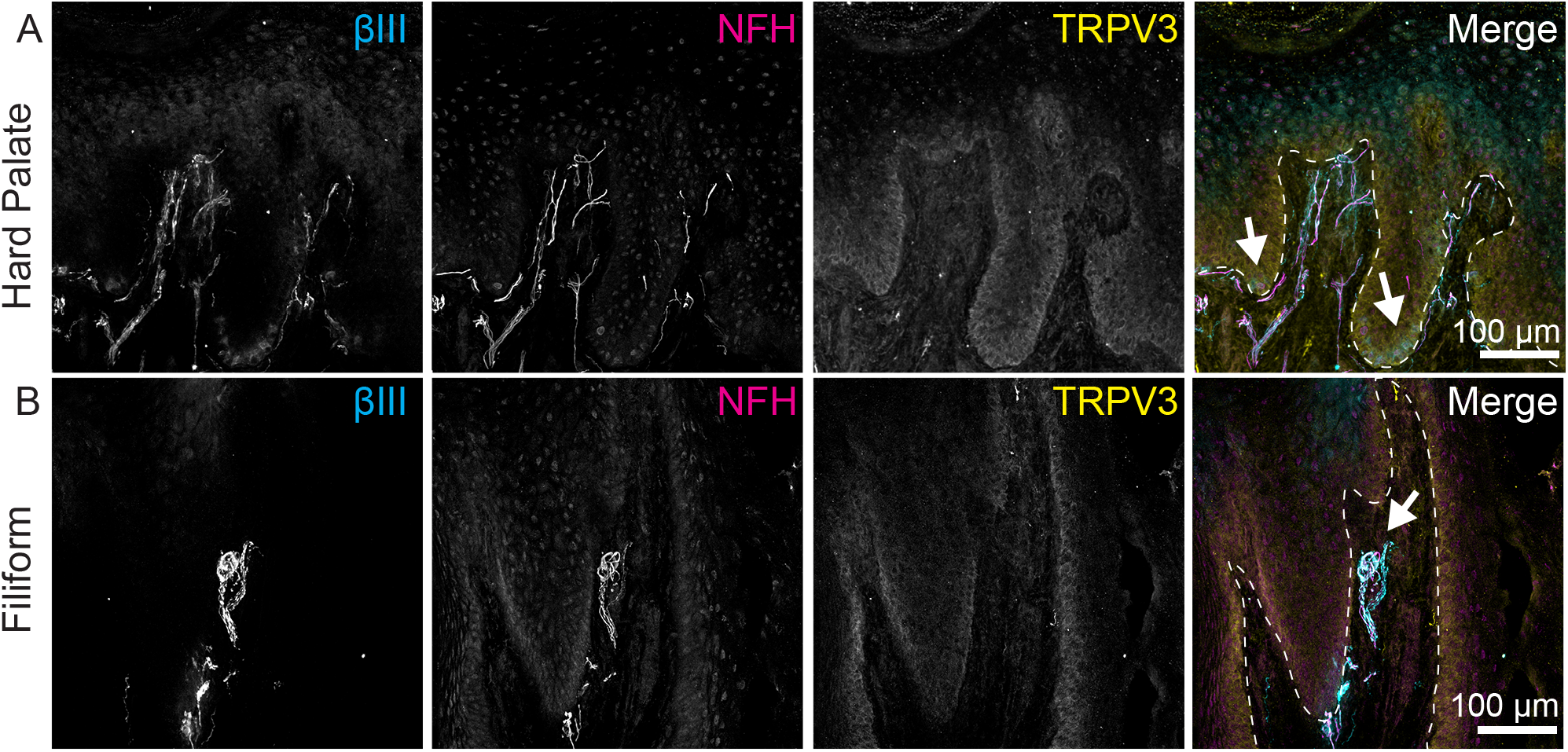
TRPV3 is expressed in basal layers of oral epithelium. **Left** column, Tuj1 anti-βIII tubulin (all afferent neurons); **2nd column**, anti-NFH antibodies (myelinated neurons); **3rd column**, Anti-TRPV3 antibody; **Right column**, Merge with TRP immunoreactivity in yellow, βIII immunoreactivity in cyan, NFH immunoreactivity in magenta. Dashed line indicates epithelia-lamina propria border. **A**. TRPV3 expression was found in basal epithelial layers of the hard palate as well as in the lamina propria. White arrows denotes areas where Merkel-cells typically concentrate (Moayedi et al. 2021). Note that Merkel cell afferents are visible with βIII tubulin staining in cyan. **B**. TRPV3 expression was found in basal epithelium of filiform papilla. A TRPV3-end bulb of Krause (arrow) is found within the lamina propria of the filiform papilla visualized with expression of βIII tubulin and NFH.

TRPV4 antibodies showed strong immunoreactivity in both tongue and hard palate (**Figure 2**). Immunoreactivity was localized in outer epithelium of hard palate mucosa and did not overlap with Merkel cells (**Figures 2A**). In the tongue, TRPV4 immunoreactivity was also identified in outer epithelial layers (**Figures 2B-C**); however, it was specifically excluded from taste buds (**Figure 2B**). Neurons innervating epithelial mucosa surrounding the taste buds, including NFH+ and NFH-afferents, extended into TRPV4+ epithelial layers. In filiform papilla, some intraepithelial nerve fibers also extend into TRPV4+ lamina (**Figure 2C**).

**Figure 2.**
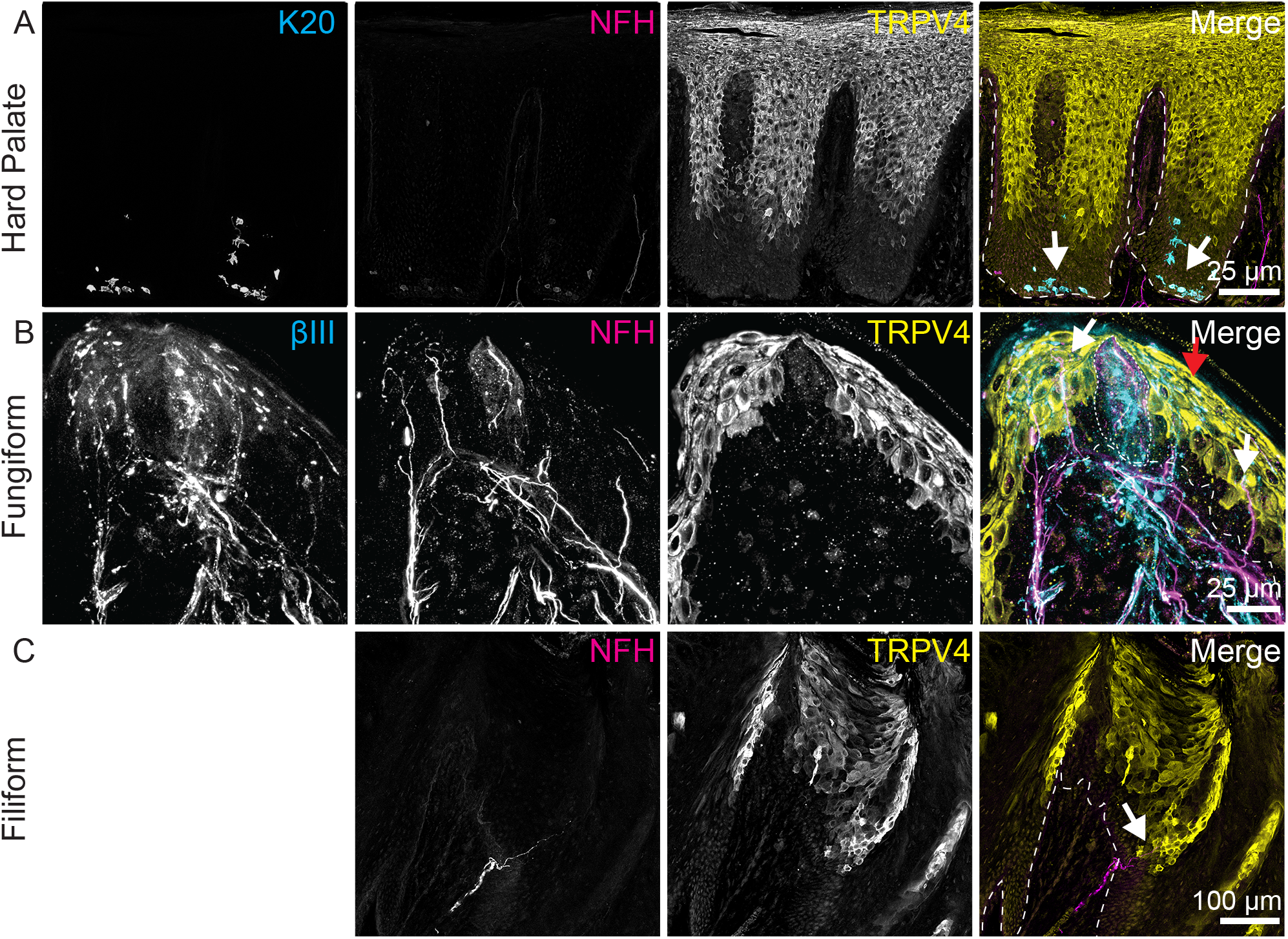
TRPV4 is expressed in apical keratinocytes. **Left** column, anti-K20 (Merkel cells) or Tuj1 anti-βIII tubulin (all afferent neurons); **2nd column**, anti-NFH antibody (myelinated neurons); **3rd column**, Anti-TRPV4 antibody; **Right column**, Merge with TRP immunoreactivity in yellow, βIII or K20 immunoreactivity in cyan, NFH immunoreactivity in magenta. Dashed line indicates epithelia-lamina propria border. Dotted lines indicate location of taste bud. **A**. Expression of TRPV4 was found in the upper layers of the hard palate epithelium. TRPV4 does not appear to be expressed in Merkel cells (white arrows) **B**. TRPV4 is expressed in apical keratinocytes in fungiform papillae epithelium, but is excluded from taste bud (white dotted line). Several nearby NFH+ and Tuj1+ neuronal afferents extend into epithelial layers expressing TRPV4 (white arrows). **C**. TRPV4 is expressed in apical epithelial cells of filiform papillae. An intraepithelial NFH+ fiber (white arrow) in close association with TRPV4+ epithelial cells is shown.

TRPA1 antibodies showed robust immunolocalization in the hard palate and tongue, with the highest density in lamina propria cells (**Figure 3**). Antigen blocking experiments confirmed specificity of TRPA1 antibodies to the antigen (**Supplementary figure 2**). Sparse TRPA1+ epithelial cells were also found with dendritic morphologies consistent with immune cells (**Figures 3A, C white arrows**). Co-staining with an antibody against CD45, a marker for immune cells, revealed that many TRPA1+ lamina propria and epithelial cells were CD45+ (**Supplementary figure 3**). To test whether neuronal processes were also TRPA1+, we analyzed 1-µm optical planes for co-labelling of TRPA1 and βIII-tubulin, a general marker for peripheral neurons. This method identified neuronal afferents that were TRPA1+ in the lamina propria of hard palate (**Figure 3B, white arrows**). In the fungiform papillae, we found TRPA1+ nerve fibers in the plexus below the taste bud (**Figure 3D, white arrows**). Interestingly, neurons innervating the taste bud were not labelled with TRPA1 antibody (**Figure 3D, purple arrows**). We next analyzed end bulbs of Krause in the lamina propria of filiform papillae (**Figure 3E-F, white arrow**). TRPA1 immunoreactivity was present in some NFH-fibers of the end bulb of Krause (**Figure 3F, white arrow**). Large neuronal fibers that were not immunoreactive to TRPA1 antibodies also contributed to the end bulb of Krause (**Figure 3F, red arrow**). Collectively, these data show that TRPA1 immunoreactivity is present in oral lamina propria cells, immune cells, and some subsets of neurons.

**Figure 3.**
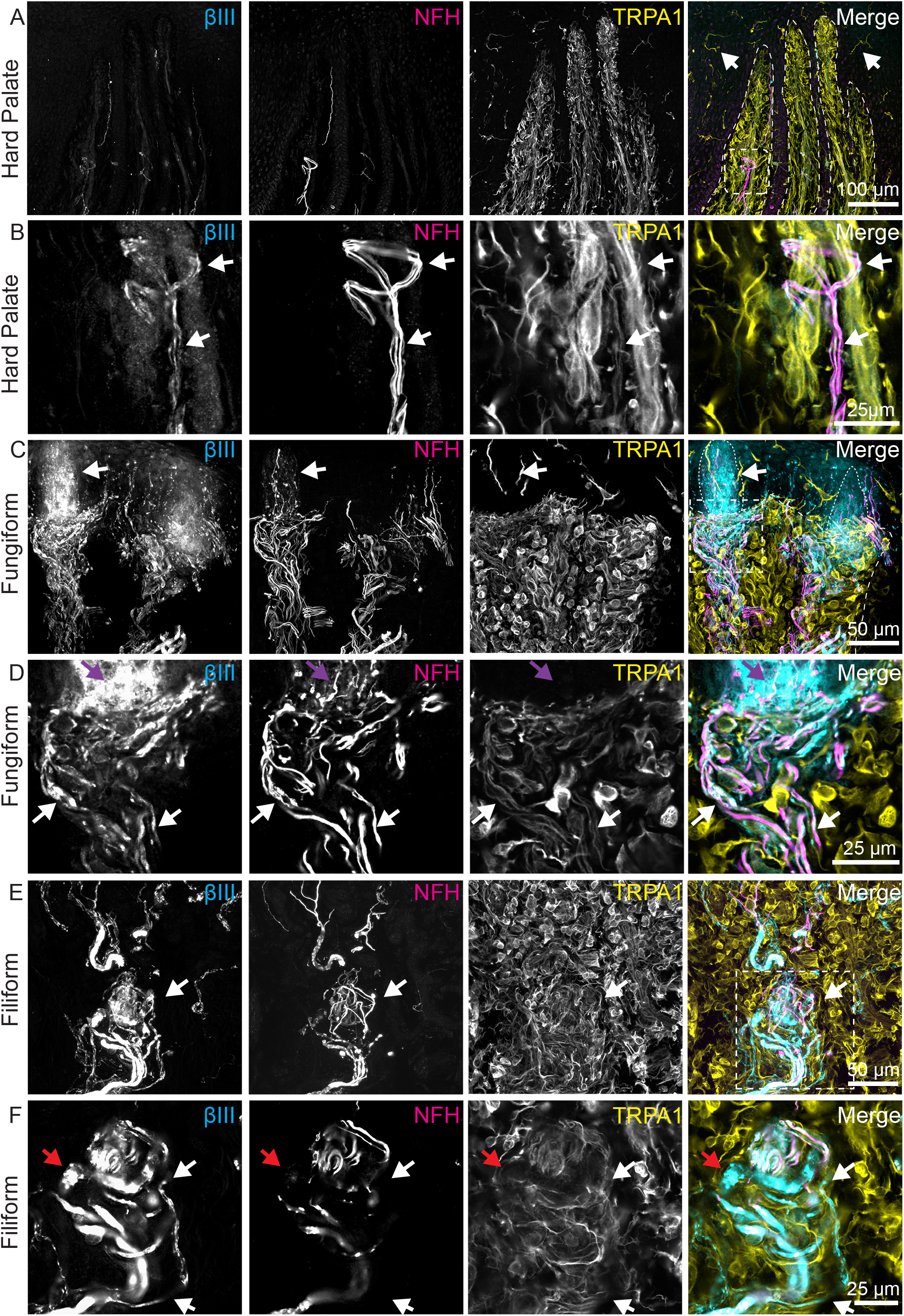
TRPA1 immunoreactivity is found in lamina propria, epithelium, and in some neuronal afferents. **Left** column, Tuj1 anti-βIII tubulin (all afferent neurons); **2nd column**, anti-NFH antibodies (myelinated neurons); **3rd column**, Anti-TRPA1 antibody; **Right column**, Merge with TRP immunoreactivity in yellow, βIII immunoreactivity in cyan, NFH immunoreactivity in magenta. Dashed line indicates epithelia-lamina propria border. Dotted lines indicate location of taste buds. Dashed box indicates area of detailed, 1 µm thick images (**B, D, F)**. **A**. TRPA1 was broadly expressed throughout lamina propria cells of the hard palate as well as sparsely in the epithelium (white arrows). A bundle of NFH+ and NFH- neuronal fibers was identified in this peg (white dashed box). White dotted box shows region in **B**. **B**. A higher magnification, single optical plane (1 µm) of the neuronal fibers in **A** showed TRPA1 expression overlapping with neurons (white arrow). **C**. TRPA1 was broadly expressed throughout lamina propria cells in fungiform papillae as well as in some epithelial cells near taste buds (white arrows). White dotted box shows region in **D**. **D**. A single optical plane (1 µm) from **C** is shown. TRPA1 expressing fibers overlap with Tuj1+ neuronal fibers (white arrows). Tuj1+ neurons that extend into epithelium do not express TRPA1 (purple arrows). **E**. TRPA1 was broadly expressed throughout lamina propria cells in filiform papillae and in end bulb of Krause (white arrow). White dashed box shows region in **F**. **F**. Single optical plane (1 µm) from **E** showed neuronal afferents expressing TRPA1 (white arrows). Larger afferents appeared to be negative for TRPA1 (red arrow).

TRPM8 immunoreactivity was next analyzed (**Figure 4**). This antibody showed broad, low-level immunoreactivity throughout epithelial and lamina propria cells with higher signal in some neurons and lamina propria cells. In the hard palate, TrpM8 immunoreactivity was widespread in lamina propria cells (**Figure 4A**). Within palate rugae, we identified TRPM8+ neurons (**Figure 4A, white arrows**) as well as TRPM8-neurons (**Figure 4A, red arrows**). In fungiform papillae (**Figure 4B**), diffuse TRPM8 immunoreactivity in the mucosa and cells of the lamina propria was found. TRPM8 immunoreactivity concentrated in the taste bud region, near the taste pore (**Figure 4B, purple arrow**). Within fungiform papillae, lamina propria, neuronal bundles were identified that with TRPM8 immunoreactivity (**Figure 4B, white arrow**), as well as TRPM8-neurons (**Figure 4B, red arrow**). TRPM8 antibody showed similar localization in tongue filiform papillae, with broad immunoreactivity in the epithelium and lamina propria, as well as in some neuronal bundles (**Figure 4C**). Within end bulbs of Krause (**Figure 4C, purple arrows)**, TRPM8 immunoreactivity overlapped with neuronal fibers, consistent with speculation that these structures might be cold receptors (Ham 1950). We also identified neuronal fibers within the lamina propria both leading to end bulbs of Krause, and unassociated with end bulbs of Krause with TRPM8 immunoreactivity (**Figure 4C, white arrows)**.

**Figure 4.**
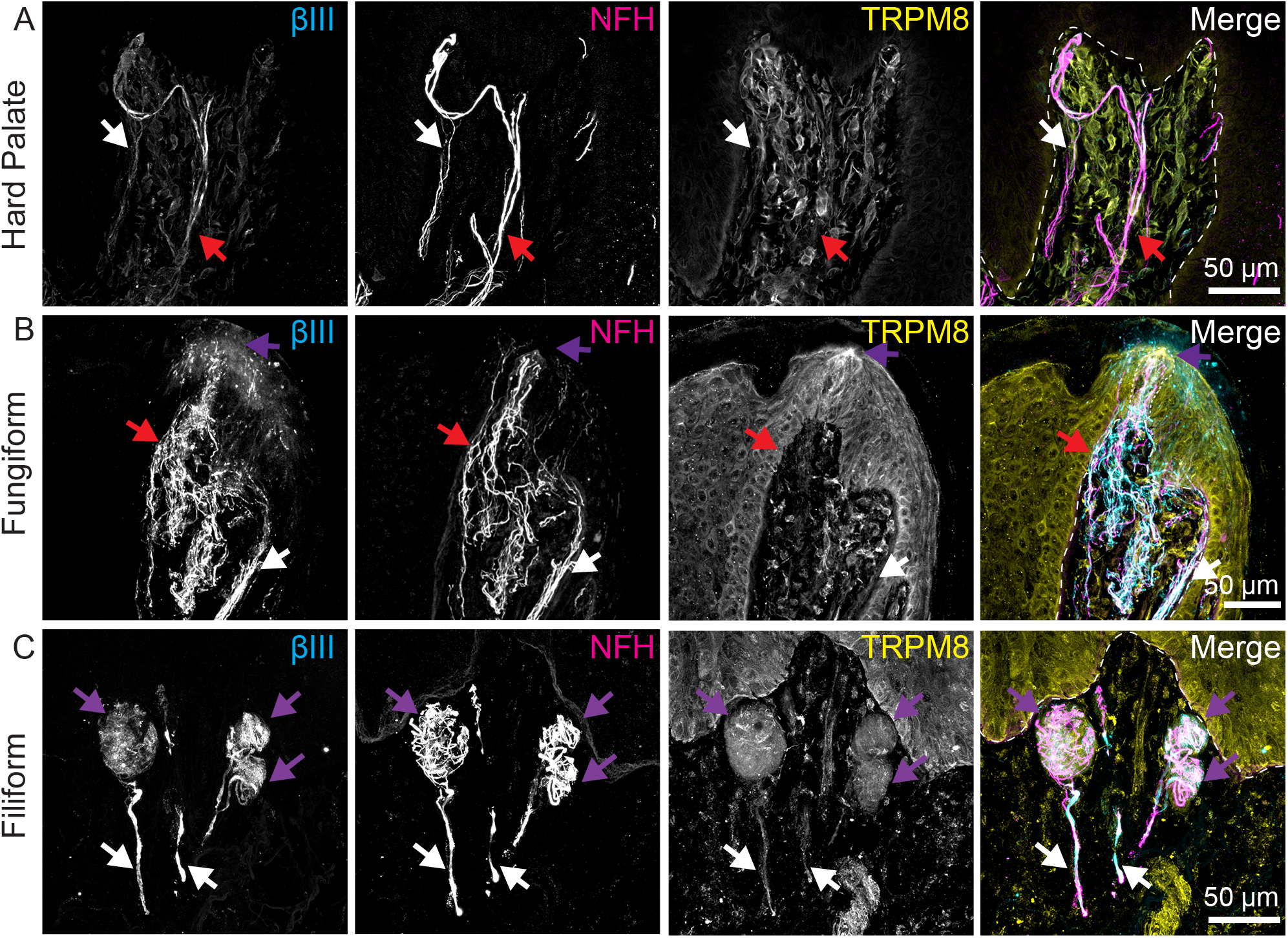
TRPM8 immunoreactivity is found throughout lamina propria, epithelium, and in neurons in oral epithelia. **Left** column, Tuj1 anti-βIII tubulin (all afferent neurons); **2nd column**, anti-NFH antibodies (myelinated neurons); **3rd column**, Anti-TRPM8 antibody; **Right column**, Merge with TRP immunoreactivity in yellow, βIII immunoreactivity in cyan, NFH immunoreactivity in magenta. Dashed line indicates epithelia-lamina propria border. Dotted line indicates location of taste bud. **A**. In hard palate epithelium, TRPM8 immunoreactivity is found in lamina propria cells and some neurons. White arrow denotes TRPM8 expression in some neurons. Red arrow denotes TRPM8 negative neurons. **B**. In tongue mucosa, TRPM8 immunoreactivity is found in epithelium and lamina propria cells. Within taste bud, a higher concentration of TRPM8 is found near the taste pore (purple arrow). Some neuronal afferents express TRPM8 (white arrow), while others are TRPM8 negative (red arrow). **C**. In filiform papillae, TRPM8 is expressed in epithelium and lamina propria cells. Purple arrows indicate end bulbs of Krause with expression of TRPM8 in some neurons within bulbs. White arrows denotes TRPM8+ neuronal afferent leading into end bulb of Krause.

Lastly, we analyzed immunolocalization of TRPV1 (**Figure 5**). In the hard palate, TRPV1+ neurons extended into the lamina propria pits of epithelial pegs (**Figure 5A, white arrows**). NFH+, TRPV1-afferents were found nearby (**Figure 5A, red arrows**). In fungiform papillae, TRPV1 expression was identified in intragemmal fibers of the taste bud, but not in nearby extragemmal fibers (**Figure 5B, white and red arrows**). In filiform papillae, TrpV1+ intraepidermal nerve fibers were found (**Figure 5C, white arrows**). Nearby, NFH+, TRPV1-neurons were also present (**Figure 5C, red arrows**). Diffuse TRPV1 immunoreactivity was identified in filiform papillae epithelium. Within the lamina propria of filiform papillae TRPV1+ fibers were frequently present (**Figure 5D, purple arrows**). We identified end bulbs of Krause in filiform papilla with high densities of NFH+ neurons; these were largely absent of TRPV1+ neurons (**Figure 5D, red arrows**). Interestingly, we found some end bulbs that had TRPV1+ neuronal immunoreactivity but with comparatively few NFH+ neurons (**Figure 5D, purple arrows**). This suggests that there is heterogeneity in neuronal composition of end bulbs of Krause.

**Figure 5.**
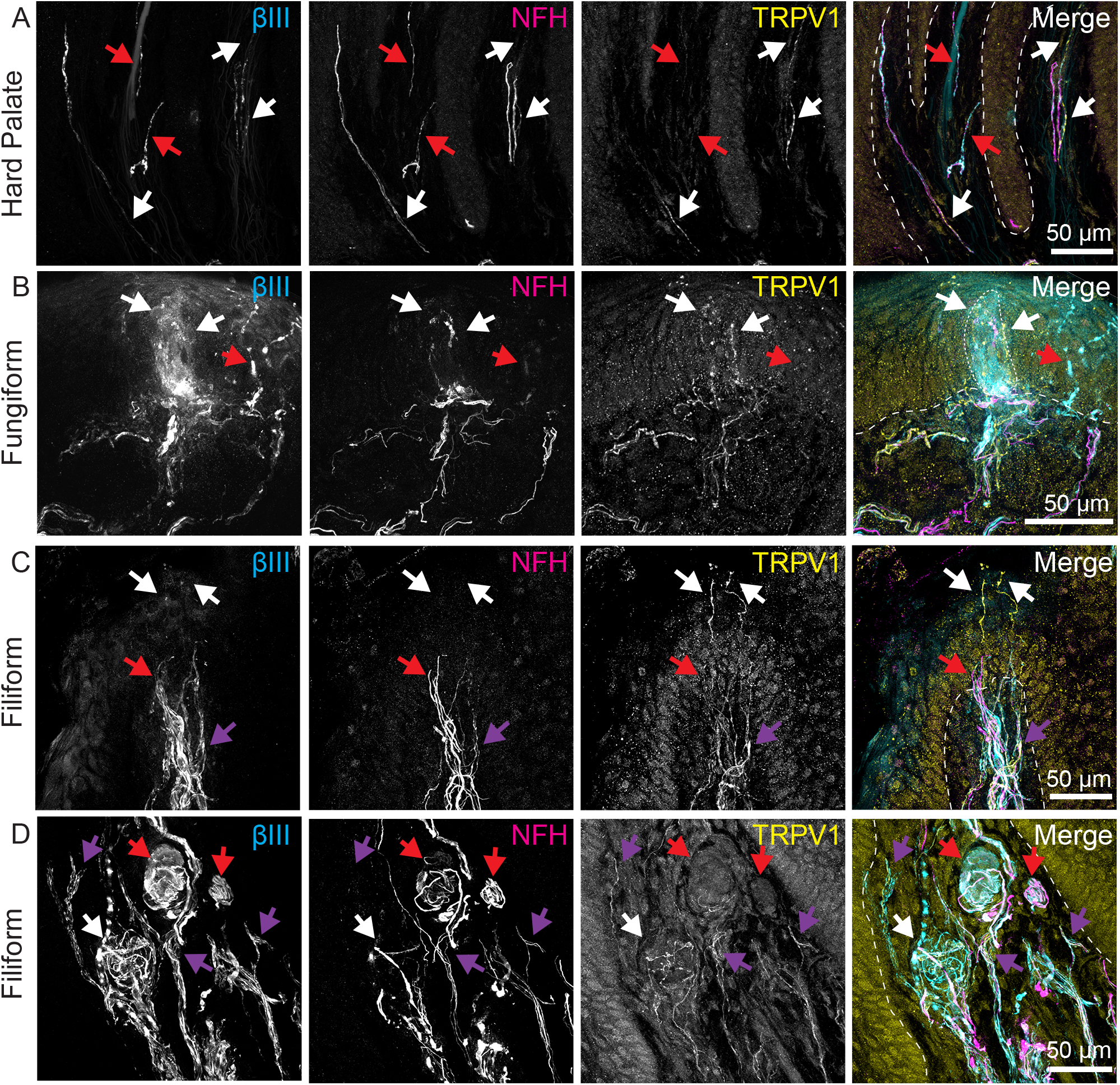
TRPV1 immunoreactivity is present in neuronal subsets and epithelium of oral mucosa. **Left** column, Tuj1 anti-βIII tubulin (all afferent neurons); **2nd column**, anti-NFH antibodies (myelinated neurons); **3rd column**, Anti-TRPV1 antibody; **Right column**, Merge with TRP immunoreactivity in yellow, βIII immunoreactivity in cyan, NFH immunoreactivity in magenta. Dashed line indicates epithelia-lamina propria border. Palate rugae from an independent sample showed TRPV1+ (blue arrows) and TRPV1-neurons (white arrows). Red box shows region in **K**. **A**. TRPV1 immunoreactivity was found in some neurons in the hard palate (white arrows). Nearby NFH+ TRPV1-neurons (red arrows) were also found. **B**. TRPV1 was identified in intragemmal fibers in fungiform papillae (white arrows). Nearby extragemmal fibers were negative for TRPV1 immunoreactivity (red arrow). **C**. TRPV1 was found intraepidermal nerve fibers in apical tips of filiform papillae (white arrows), as well as bundle of TRPV1 fiber in the lamina propria (purple arrow). Nearby NFH+ TRPV1-fibers were also found (red arrow). Light TRPV1 immunoreactivity was found in epithelial cells. **D**. End bulbs of Krause were found in lamina propria of filiform papilla. TRPV1+ fibers were found throughout lamina propria (purple arrows). One end bulb of Krause in this bundle had a low density of NFH+ fibers and high density of TRPV1+ fibers (white arrow). Two additional end bulbs of Krause had a high density of NFH+ fibers and low density of TRPV1+ fibers (red arrows).

## Discussion

TRP channels are widely expressed in the oral cavity and subserve a variety of functions including taste transduction, somatosensation and stress responses. Despite this, expression of TRP channels in oral mucosa are not well defined. In this work, we describe expression of somatosensory TRP channels in tongue and hard palate (**Supplementary Figure 4)**. We identified varied patterns of expression in epithelium, lamina propria and neuronal afferents.

Oral epithelial cells display rapid turnover and fast wound healing rates due to intrinsic differences in oral stem cells and keratinocytes (Andl et al. 2016; Iglesias-Bartolome et al. 2018). During homeostatic turnover and wound healing, epithelial barrier integrity must be maintained to prevent infection from oral bacterium; thus, molecules involved in barrier integrity and epithelial maintenance are particularly important in oral epithelium. TRPV3 and TRPV4 are warm activated channels expressed in keratinocytes and have essential roles in epidermal development and homeostasis (Chung et al. 2004b). TRPV3 has been shown to be important for keratinocyte development and hair morphogenesis while TRPV4 is essential for skin barrier formation (Blaydon and Kelsell 2014; Cheng et al. 2010; Sokabe et al. 2010). In the oral cavity, TRPV3 and TRPV4 were expressed primarily in epithelium with inverse distributions. TRPV3 immunoreactivity was observed throughout basal layers of epithelium in both the tongue and hard palate, similar to previous findings (Xu et al. 2006). In contrast, TRPV4 immunoreactivity was found in apical layers of oral cavity epithelium, and was excluded from taste cells and Merkel cells. This expression pattern in outer keratinocyte layers in oral tissues is consistent with expression in palmar keratinocytes (Blaydon and Kelsell 2014). The expression of TRPV3 and TRPV4 was consistent with previous studies showing that oral epithelia respond to TRPV3 and TRPV4 agonists including camphor, 4α-phorbol-12,13 didecanoate (4α-PDD), 2-aminoethoxydiphenyl borate (2-APB) (Wang et al. 2011). Based on expression patterns and known functions of TRP channels, we can build hypotheses on the functions in oral tissues. TRPV3 likely plays an important role in oral epithelial growth and renewal, particularly after damage (Aijima et al. 2015). TRPV4 is likely playing a role in oral barrier formation and may mediate inflammatory signaling and pain after tissue damage (Moore et al. 2013; Rajasekhar et al. 2017; Sokabe et al. 2010). Furthermore, as TRPV4 responds to shear stress, it may also function in cell signaling in response to epithelial stretch (Rajasekhar et al. 2017).

TRPA1 is a key damage sensor in many organs and tissues (Talavera et al. 2020). Consistent with this, TRPA1 immunoreactivity was identified predominantly in cell types that are poised report tissue injury including cells in the lamina propria and immune cells. The widespread expression of TRPA1 in the lamina propria of hard palate and tongue suggests that it is expressed broadly in oral fibroblasts. TRPA1 is functionally expressed in human dental fibroblasts, suggesting that this is a conserved TRPA1 pattern of expression in the oral cavity (El Karim et al. 2011). TRPA1 expression within oral lamina propria provides an optimal localization to play a role in remodeling due to tissue damage. Subpopulations of TRPA1+ cells in lamina propria and epithelium expressed CD45, a well-established marker of immune cells, indicating that TRPA1+ is widely expressing in immune cells of oral mucosa. The widespread expression of TRPA1 is consistent with a role of TRPA1 in inflammation in oral cavity. Expression in immune cells and fibroblast positions this channel to signal the presence of noxious compounds or tissue damage.

In addition to roles in responding to cellular damage, TRPA1 is activated by pungent compounds, like wasabi and allicin, and plays a role in chemesthesis during flavor construction (Talavera et al. 2020). TRPA1 immunoreactivity was observed in afferents innervating both the tongue and hard palate, largely excluding large diameter neuronal endings. Expression in neuronal afferents confers these neuronal endings with the ability to detect pungent compounds in the mouth, or to take part in pain, itch, and thermal signal transduction.

Temperature sensation in oral tissues is an important aspect of flavor construction. TRPM8 is a cold and menthol-activated receptor that is essential for cold and warm sensations (Moore et al. 2018; Paricio-Montesinos et al. 2020). Neuronal immunoreactivity to TRPM8 was identified in hard palate and tongue mucosa. In the filiform papillae of the tongue, TRPM8+ neurons were also found within end bulbs of Krause, indicating that these structures may include cold sensitive afferents as initially theorized (Ham 1950). TRPM8 immunoreactivity was also found to concentrate around the taste bud in fungiform papillae of the tongue. Expression around fungiform papilla taste buds is likely to contribute to chemesthesis of flavors of compounds like menthol and eucalyptol as well as transduction of cold sensations. In addition to neuronal localization, we identified TRPM8 immunoreactivity in oral fibroblasts, consistent with previous findings in human oral tissues (El Karim et al. 2011). Here, TRPM8 may take part in remodeling due to chemical or thermal activation.

TRPV1 is expressed in nociceptive afferents and epithelial cells and is responsive to capsaicin heat, pH, and histamine, amongst other compounds (Moore et al. 2018). TRPV1 activating compounds are particularly important in regards to oral function as they mediate flavor construction, oral homeostasis, and response to pathogens and injury. We found light TRPV1 immunoreactivity in the epithelium of tongue and hard palate. Epithelial expression of TRPV1 has been shown previously in skin, where it plays a role in keratinocyte migration, and epidermal barrier integrity (Blaydon and Kelsell 2014). Oral epithelia respond to capsaicin, suggesting that TRPV1 expression is functional in oral keratinocytes (Wang et al. 2011).

Neuronal TRPV1 immunoreactivity was widespread in the oral cavity. We identified TRPV1+ free nerve endings in the tongue and hard palate, including intraepithelial nerve fibers. These neurons would be suspected to take part in nociception, temperature sensation, and chemesthesis. In fungiform taste buds, we identified TRPV1 immunoreactivity in intragemmal fibers of the taste bud, but not in extragemmal fibers. These intragemmal fibers are ideally positioned to take part in flavor construction by transducing temperature and spiciness of foods. Within filiform papillae of the tongue, TRPV1 staining was surprisingly identified in neurons associated with end bulbs of Krause. These findings suggest a previously unappreciated diversity in the population of neurons within end bulbs of Krause, having both myelinated, likely mechanosensory neurons intermingled with unmyelinated TRPV1+ and TRPM8+ populations that may take part in thermosensation. We also noted diversity in end bulbs of Krause compositions within a single filiform papilla, with some having dense NFH+ myelinated afferents while others have fewer myelinated afferents and higher unmyelinated, TRPV1+ afferents. The finding of independent neuronal afferent subtypes within a single corpuscle are reminiscent of findings in Meissner’s corpuscles, where two distinct populations of mechanoreceptors have been found in mice, and where both myelinated and unmyelinated neurons have been identified in humans (Cauna 1956; Neubarth et al. 2020). Future studies are required to parse out differences in end bulbs of Krause populations and to better understand the functions of these structures in somatosensation.

TRPV1 expressing neurons play important roles in nociception, inflammatory pain, and neuropathic pain (Moore et al. 2018). Patients with burning mouth syndrome (BMS) have an increase in both the presence of TRPV1+ nerve fibers and in epithelial TRPV1 expression (Borsani et al. 2014; Yilmaz et al. 2007). This suggests that expression of TRPV1 could be linked to the pathogenesis of BMS. Future studies should be performed investigating whether TRPV1 expression is upregulated in particular ending types, such as intraepidermal nerve fibers or end bulbs of Krause, in BMS patients.

There are important limitations to consider when interpreting findings from this study. Antibody immunoreactivity might not recapitulate the protein expression pattern due to cross-reactivity with other epitopes. Future studies should compare results with antibodies against other protein epitopes, analyze RNA expression, and test functional expression of TRP channels in oral tissues. A second limitation is that tissue donors for tongue biopsies were primarily female. Future studies should analyze sex differences in TRP channel expression.

In summary, we describe the immunohistochemical localization of TRP channels throughout human healthy oral epithelium. Oral TRP channels are poised to take part in a myriad of functions including epithelial integrity, epithelial development, response to injury, thermoception, nociception, and flavor construction. Future studies are needed to parse out roles for TRP channels in oral pathologies.

## Acknowledgments

We thank Benjamin Le Reverend for discussions during the conceptualization of this project, Rong Du for assistance in providing human foreskin samples, and Joy Chen for assistance with oral biopsy collection. This work was supported by SPN, SA; Berrie Foundation Initiative on the Neurobiology of Obesity (EAL, YM); Thompson Family Foundation Initiative on Chemotherapy Induced Peripheral Neuropathy and Sensory Neuroscience (YM); and NIH NIAMS R01AR051219 (EAL). Imaging was performed with support from the Zuckerman Institute’s Cellular Imaging Platform. Human foreskin samples were collected with support from the Columbia University Skin Disease Resource Based Center (epiCURE, P30AR069632).

