## Supplemental Figures for "Localization of TRP channels in healthy oral mucosa from human donors"

**Supplementary Table 1. Collected biopsies**

| Site of Biopsy | Age | Gender |
| --- | --- | --- |
| Palate Rugae | 40 | F |
| Palate Rugae | 30 | F |
| Palate Rugae | 43 | F |
| Palate Rugae | 28 | M |
| Palate Rugae | 35 | M |
| Tongue Back | 45 | F |
| Tongue Front | 38 | F |
| Tongue Front | 43 | F |
| Tongue Front | 33 | F |
| Tongue Front | 27 | F |
| Tongue Front | 28 | F |
| Tongue Front | 28 | M |

**Supplementary table 2. Antibodies used in this study**

| Antibody | Supplier | Catalog # | Lot | Dilution | RRID |
|---|---|---|---|---|---|
| Mouse anti-Keratin 20 | Abcam | Ab854 | GR157163-5 | 1:100 | AB_2133708 |
| Chicken anti-Neurofilament-Heavy | Abcam | Ab4680 | GR310109-11 | 1:5000 | AB_304560 |
| Mouse anti-βIII tubulin (Tuj1) | Neuromics | MO15013 | 402154 & 402360 | 1:100 | AB_2737114 |
| Moue anti-CD45 | Abcam | Ab781 | GR3233952-8 | 1:100 | AB_306098 |
| Rabbit anti-TRPA1 | Alomone labs | ACC-037 | ACC037AN17 | 1:500 | AB_2040232 |
| Rabbit anti-TRPV1 | Abcam | Ab3487 | GR3219961-4 | 1:500 | AB_2209009 |
| Rabbit anti-TRPM8 | Alomone labs | ACC-049 | ACC049AN15 | 1:100 | AB_2040254 |
| Rabbit anti-TRPV4 | Lifespan Bio | LS-A8583 | 61861 | 1:100 | AB_592927 |
| Rabbit anti-TRPV3 | Alomone Labs | ACC-033 | ACC033AN02 | 1:100 | AB_2040261 |

**Supplementary Figure 1. Biopsy sites**

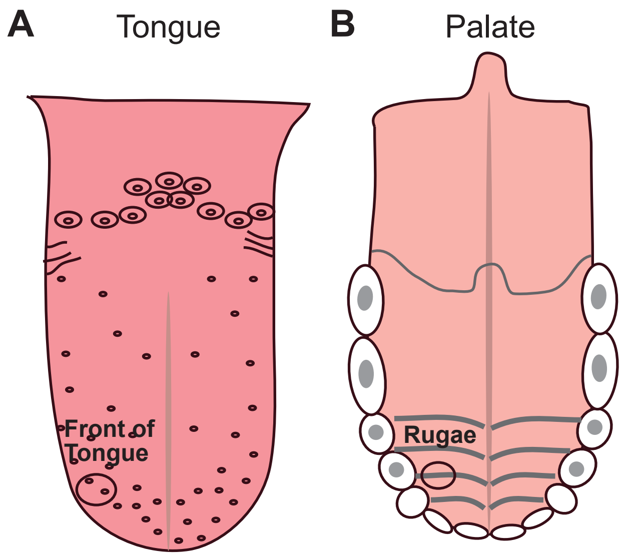


**Figure 1. Schematics of collection sites.** Schematics showing approximate collection sites on tongue and hard palate are shown. Tongue biopsies (**A**) were collected towards the front of the tongue and aimed to include a taste bud. Palate Biopsies (**B**) were collected towards that lateral aspect of the hard palate and included a portion of a rugae.

**Supplementary Figure 2. Antigen blocking**

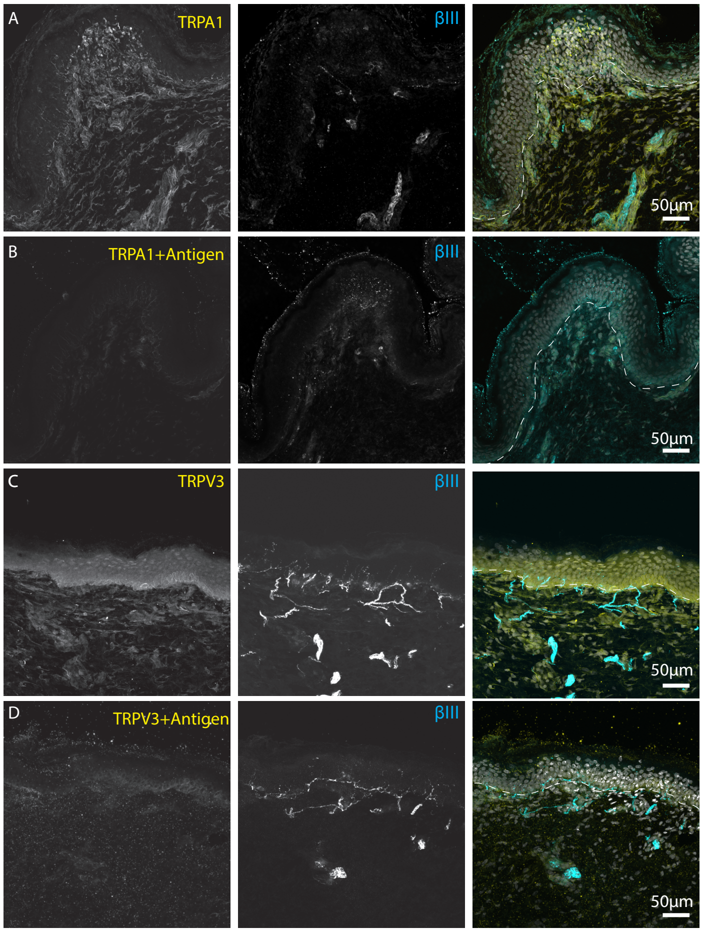


**Supplementary Figure 2. Antigen blocking shows specificity of TrpA1 and TrpV3 antibodies.**

**Left** column, Anti-TRP antibody; **Middle column,** Tuj1 anti-βIII tubulin (all afferent neurons); **Right column**: Merge with TRP immunoreactivity in yellow, βIII immunoreactivity in cyan, and DAPI to visualize nuclei in gray.

- **A.** Antibody staining of human foreskin tissue was performed using an antibody against TRPA1. Immunoreactivity was found throughout the epidermal and dermal layers of skin. Dashed line denotes epidermal-dermal boarder.
- **B.** Antigen blocking completely eliminated TRPA1 immunoreactivity in skin, indicating specificity of this antibody to the epitope.
- **C.** TRPV3 immunostaining was optimized using human skin specimens. Immunoreactivity was found throughout the epidermal and dermal layers, with particular intensity towards the basal epidermis.
- **D.** Antigen blocking reduced TRPV3 immunoreactivity, indicating specificity of this antibody for the TRPV3 epitope.

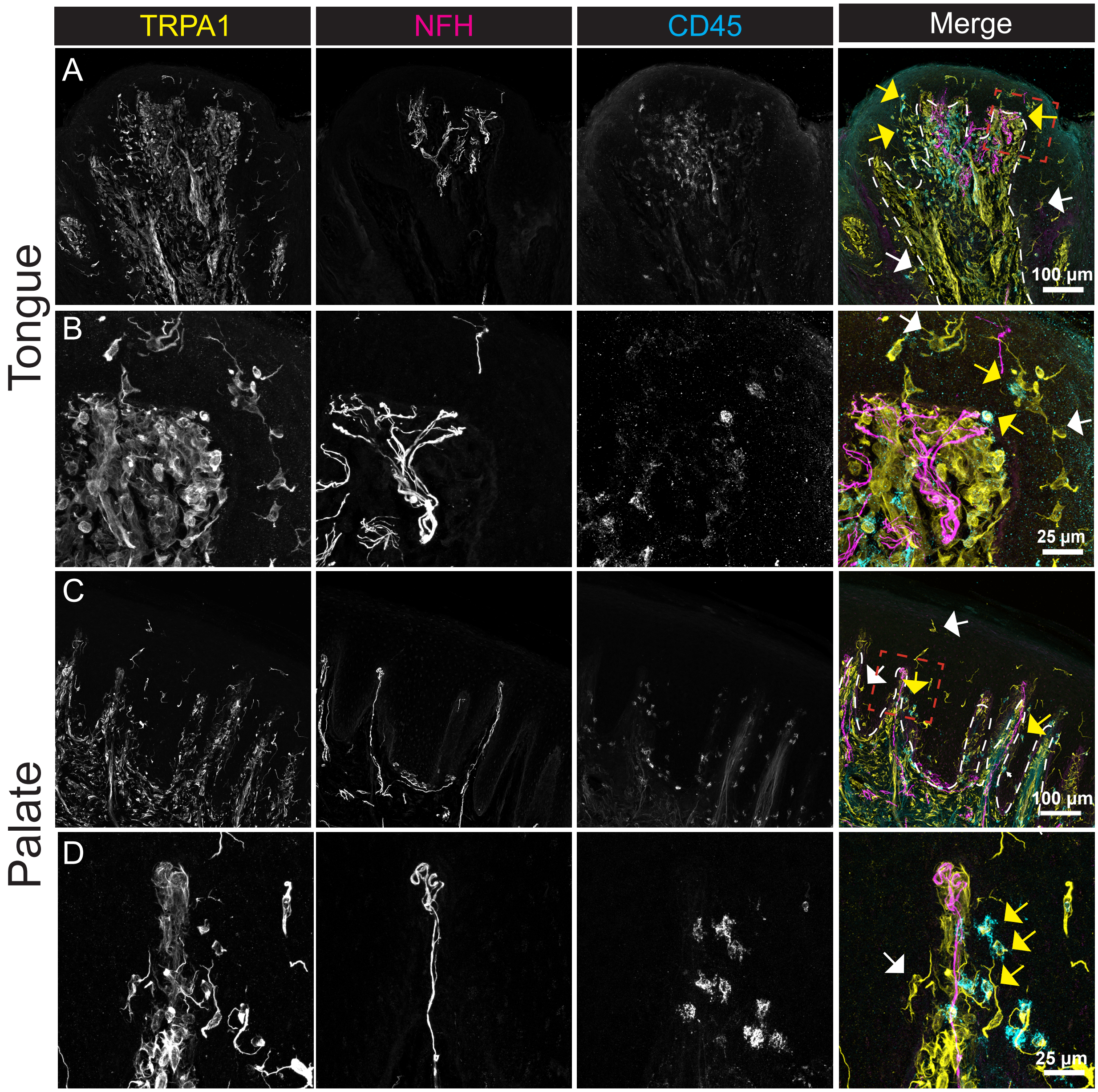


**Supplementary Figure 3. TrpA1 immunoreactivity co-expresses with CD45 in (A-C) and hard palate rugae (D-F).**

**Left** column, Anti-TRPA1 antibody; **2nd column**, NFH antibody (myelinated neurons); **3rd column**, anti-CD45 antibody (immune cells); **Right column**, Merge with TRP immunoreactivity in yellow, CD45 immunoreactivity in cyan, NFH immunoreactivity in magenta.

- **A.** TRPA1 was broadly expressed throughout lamina propria cells and some cells in epithelial layer of tongue. Co-expression of TRPA1 and CD45 was identified in some epithelial cells (yellow arrows). TrpA1+ cells were also found that did not co-express CD45 (white arrows). Red box shows region in **B.**
- **B.** A higher magnification view of **A**.

**C.** TRPA1 shows a similar pattern of expression in the hard palate as in tongue. Again, both cells that co-expressed CD45 (yellow arrows) and those that did not (white arrows) were identified.

**D.** Higher magnification view of **C**.

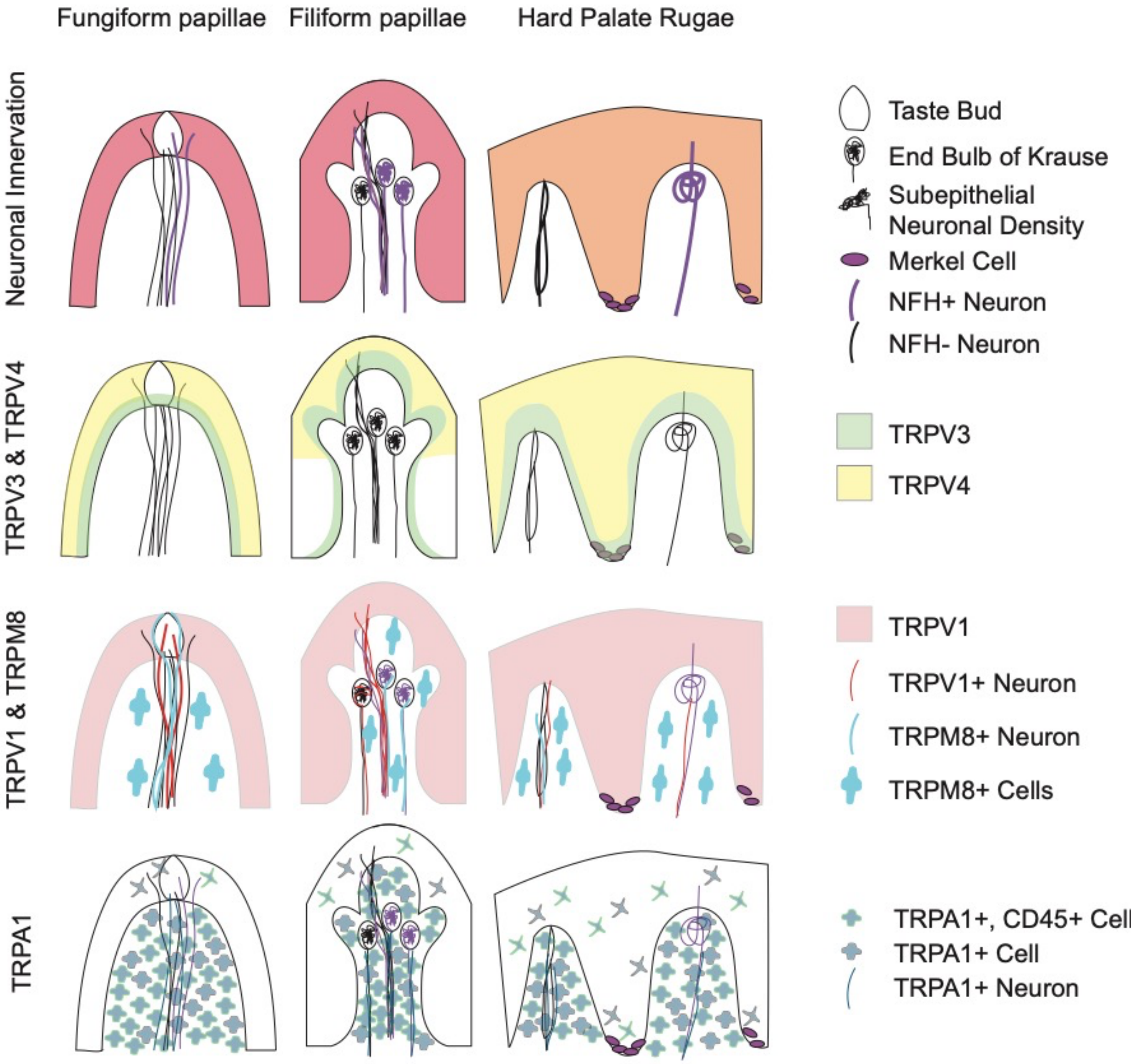


**Supplementary Figure 4. Summary of TRP channel expression in oral tissues.** Distribution of TRP immunoreactivity in tongue and hard palate samples are shown.
